# Expression of Pokeweed Antiviral Protein Isoform S1 (PAP-S1) and of Ricin-A-Chain/PAP-S1 novel fusion protein (RTA/PAP-S1) in *Escherichia coli* and their comparative inhibition of protein synthesis *in vitro*

**DOI:** 10.1101/137919

**Authors:** Yasser Hassan, Sherry Ogg

## Abstract

Fusion protein therapeutics engineering is advancing to meet the need for novel medicine. Herein, we further characterize the development of novel RTA & PAP-S1 antiviral fusion proteins. In brief, RTA/PAP-S1 and PAP-S1/RTA fusion proteins were produced in both cell free and *E. coli in vivo* expression systems, purified by His-tag affinity chromatography, and protein synthesis inhibitory activity assayed by comparison to the production of a control protein, CalmL3. Results showed that the RTA/PAP-S1 fusion protein is amenable to standardized production and purification and has both increased potency and less toxicity compared to either RTA or PAP-S1 alone. Thus, this research highlights the developmental potential of novel fusion proteins with reduced cytotoxic risk and increased potency.

## 1. Introduction

Pokeweed antiviral proteins (PAPs) are potent type I Ribosome Inactivating Proteins (RIPs) expressed in several organs of the plant pokeweed (*Phytolacca americana*), as reviewed by Domashevskiy and Goss [1]. They are chiefly secreted and bound within the plant cell wall matrix. Here, they are known to function in defense against pathogens through the inhibition of both prokaryotic and eukaryotic ribosomes and protein synthesis. Among the PAP gene family, different genes are expressed in various tissues and at different stages of development in *Phytolacca americana*. PAP, PAP II, PAP-S1, PAP-S2, and PAP-R are the forms that appear in spring leaves, summer leaves, isoform S1 and S2 in seeds, and roots, respectively. The molecular weight ranges from 29 kDa for PAP to 30 kDa for PAP-S’s [2]. PAP-S1 has been identified as the most effective in inhibiting protein synthesis *in vitro* [3]. PAPs possess antiviral activity on a wide range of plant and human viruses; different forms of PAP expressed in transgenic plants leads to broad-spectrum resistance to viral and fungal infections [4-5]. Relevant to the recent Zika epidemic, PAP is efficient against Japanese encephalitis virus [6]. Antiviral activity is also present against HIV-1 [7], human T-cell leukemia virus-1 (HTLV-1) [8], herpes simplex virus (HSV) [9], influenza [10], hepatitis B virus (HBV) [11], and poliovirus [12]. Moreover, different forms of PAP have moderate cytotoxicity to non-infected cells and, thus, offer unique opportunities for new applications in therapy and as a protective protein against pathogens in transgenic plants.

Ricin is produced in the seeds of the castor oil plant, *Ricinus communis*, and is one of the most potent type II RIPs, as reviewed by Lord et al [13]. It can efficiently deliver its A chain into the cytosol of cells through the action of its B chain. The B chain serves as a galactose/N-acetylgalactosamine binding domain (lectin) and is linked to the A chain via disulfide bonds. After the ricin B chain binds complex carbohydrates on the surface of eukaryotic cells containing either terminal N-acetylgalactosamine or beta-1,4-linked galactose residues, it is endocytosed via clathrin-dependent as well as clathrin-independent mechanisms and is thereafter delivered into the early endosomes. It is then transported to the Golgi apparatus by retrograde transport to reach the endoplasmic reticulum (ER) where its disulfide bonds are cleaved by thioredoxin reductases and disulfide isomerases. The median lethal dose (LD50) of ricin is around 22 micrograms per kilogram of body weight if the exposure is from injection or inhalation (1.78 milligram for an average adult). It is important to note that the ricin A chain (RTA) on its own has less than 0.01% of the toxicity of the native lectin in a cell culture test system. It was furthermore shown that RTA alone had no activity on non-infected and tobacco mosaic virus (TMV)-infected tobacco protoplasts alike. Though there are currently no commercially available therapeutic applications, RTA is extensively studied in the development of immunotoxins [14].

The therapeutic potential of PAP and RTA has been explored for over thirty years, though side effects have limited clinical application. As evaluated by Benigni et al [15], while these proteins have shown very low cytotoxicity to non-infected cells, PAP administration in mouse models has resulted in hepatic, renal and gastrointestinal tract damage with an LD50 as low as 1.6mg/Kg. Interestingly, RTA shows no toxicity even at high doses with similar half-life times. However, all RIP’s show immunosuppressive effects to various degrees. Other studies have described the various dose-limiting side effects of these proteins when used as immunotoxins (i.e. vascular leak syndrome, hemolytic uremic syndrome, and pluritis, among others) [16-17]. Excitingly, some patients achieved complete or partial remission against Refractory B-Lineage Acute Lymphoblastic Leukemia with sub-toxic dosages, for example.

Fusion and hybrid PAP proteins have also been developed in pursuit of selectively targeting infected cells and selectively recognizing viral components, though with limited success [18-19]. Indeed, the engineering of novel therapeutic fusion proteins with higher specificity, selectivity, and potency with fewer side effects is a leading strategy in drug development.

Thus, this research furthers study of a previously created and functional novel fusion protein between RTA and PAP-S1 [20]. Here, we describe the increased potency over PAP-S1 alone, selectivity for infected cells, and reduced side effects associated with dosage. Additionally, we describe the development of an adequate and scalable production system that enabled accurate determination of PAP-S1 and RTA/PAP-S1 protein synthesis inhibition *in vitro*.

## 2. Results

### 2.1 Production and Purification of recombinant Proteins in Cell Free Expression System

The proteins were expressed and bands visible where expected (Figure 1). It is noted that expression of the three recombinant proteins is very low compared to the control protein and, more importantly, with a much lower purity; expression of PAP-S1 is the lowest. The low levels of expression of the recombinant proteins were expected as the proteins are known to be toxic to prokaryotic ribosomes. Total protein content and purity of each sample is shown in Table 1.

**Table 1.**
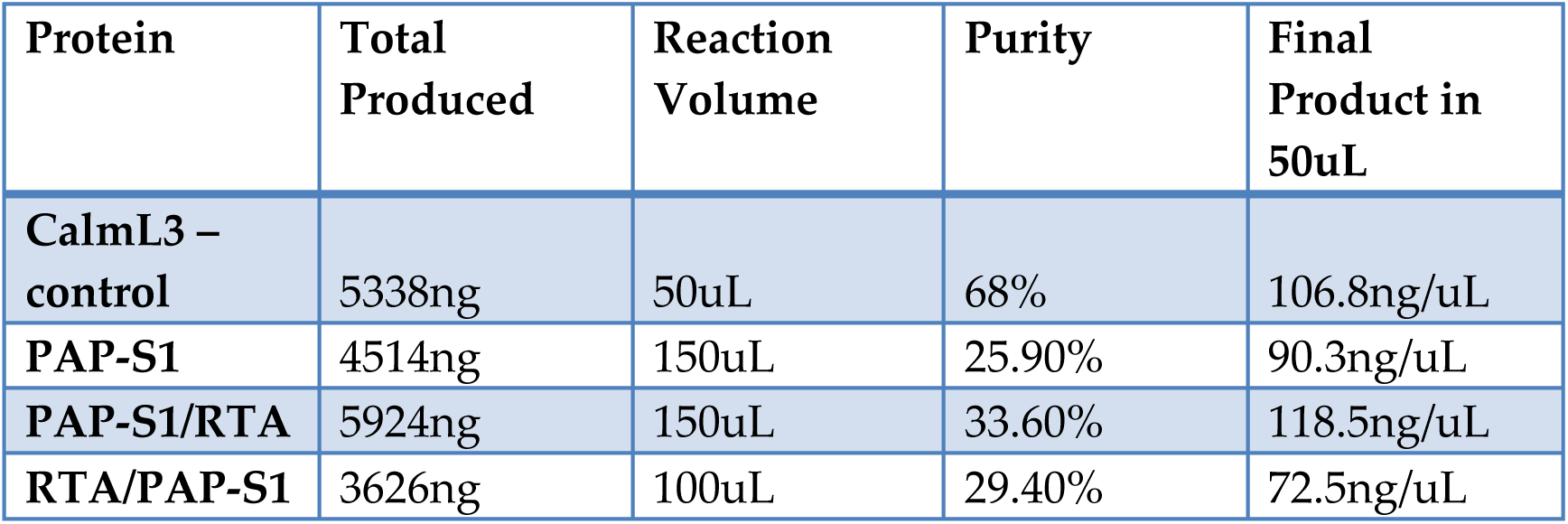
Total protein content and purity. For each sample, protein content was determined using Qubit™ 3.0 Fluorometry; purity was determined by GelAnalyzer 2010 using values from Figure 1S (See Appendix 2 for details).

**Figure 1.**
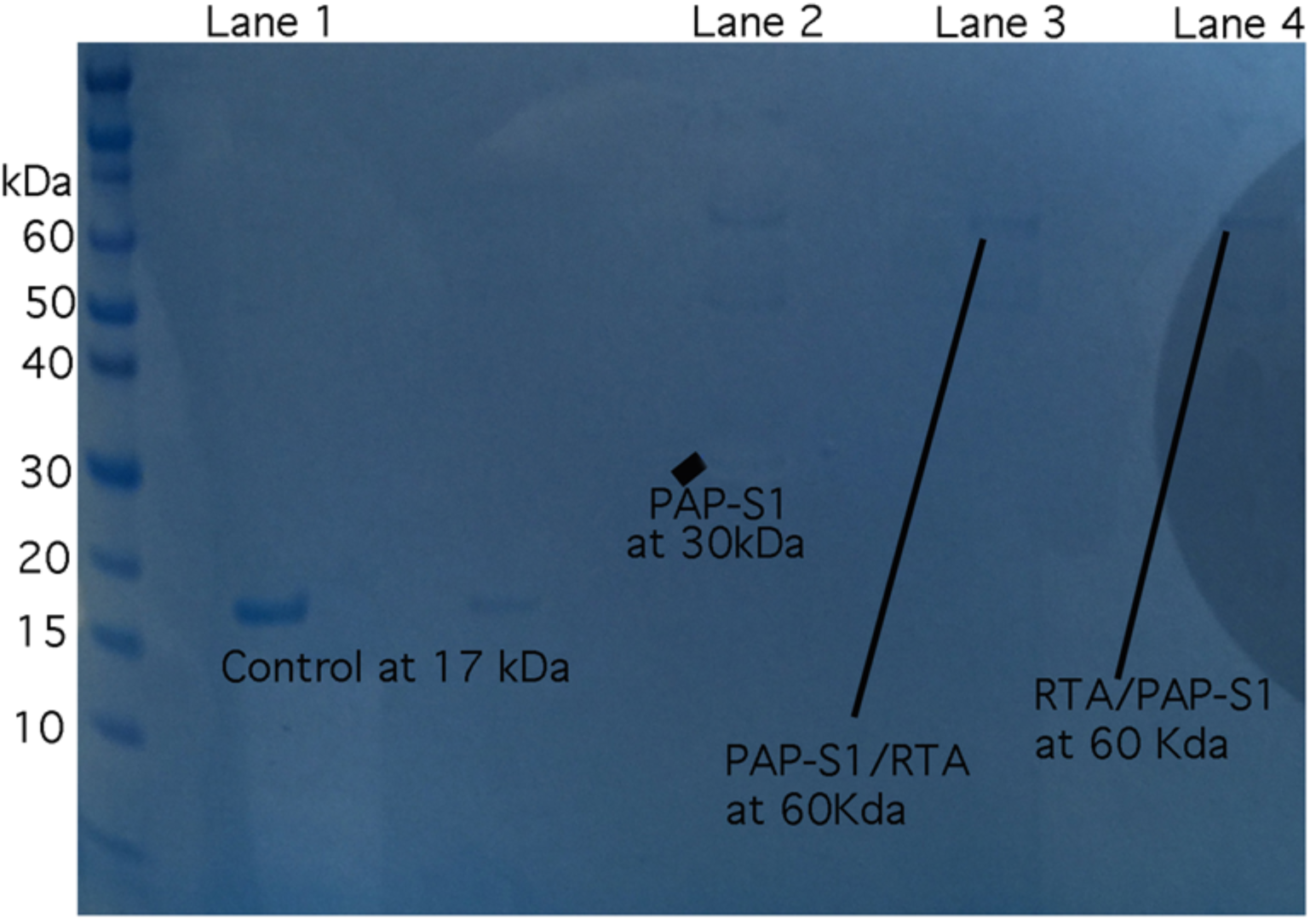
Recombinant proteins gel stained with Coomassie blue after His-Tag purification. Lane 1: Control protein after His-Tag Purification. The band is clearly visible at 17kDa. Lane 2: PAP-S1 recombinant protein at the 32kDa line. Lane 3: PAP-S1/RTA recombinant protein at the 60kDa line. Lane 4: RTA/PAP-S1 recombinant protein at the 60kDa line *All other bands are due to proteins going through the His-Tag purification column from the initial expression reaction.

### 2.2 Inhibitory activity of recombinant proteins on E. coli protein synthesis

Two different prokaryotic kits, the *E. coli* S30 T7 High-Yield Protein Expression System and the Expressway™ Mini Cell-Free Expression System, were used to control for variances in production systems. The same control protein was produced, namely CalmL3. Various concentrations of recombinant proteins were added to each reaction and the amount of CalmL3 protein produced was compared to the amount of CalmL3 produced without the addition of any recombinant protein. These values were plotted as percent protein inhibition compared to control versus concentration (Figure 2). The IC50 of PAP-S1/RTA was found to be around 460nM while the IC50 of RTA/PAP-S1 was around 241nM (based on Figure 6S and Figure 7S, see appendix 3 for details); the IC50 of PAP-S1 is known to be around 280nM [3]. Those initial results confirm RTA/PAP-S1 as more potent than PAP-S1/RTA. This was expected as the C terminal is known to play a major role in activity [23]. Those results also confirm that RTA/PAP-S1 is more potent than PAP-S1 alone. RTA/PAP-S1 will thus be the subject of the rest of the paper. The increased activity of RTA/PAP-S1 compared to PAP-S1 alone can probably be explained by the fact that PAP-S1 and RTA do not dock onto the ribosome at the same site. Indeed, it was found that after PAP-S1 partially depurinates the *E. coli* ribosome, RTA is able to depurinate the same ribosome while RTA cannot depurinate an intact *E. coli* ribosome on its own [19-26].

**Figure 2.**
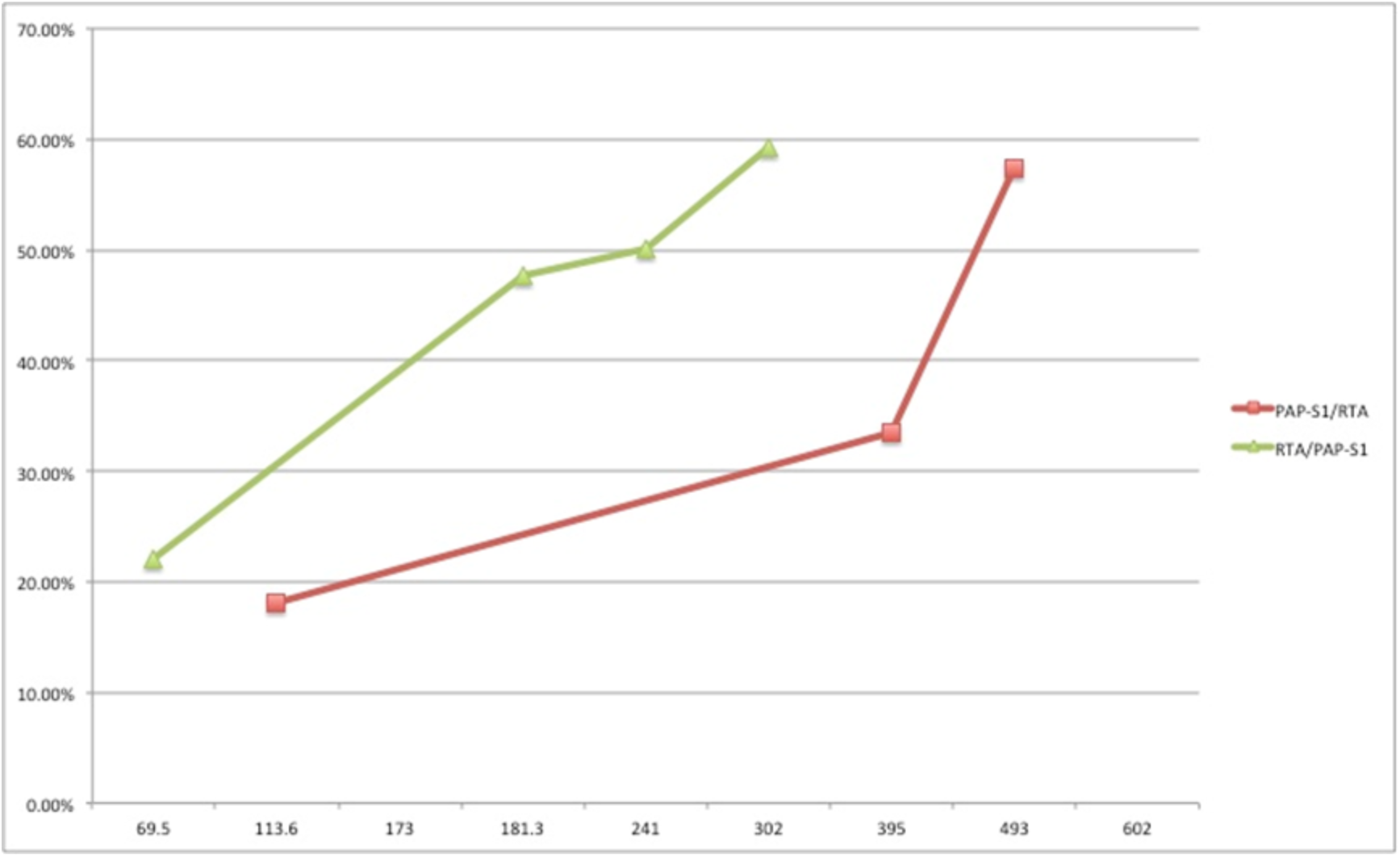
Percent protein inhibition compared to control. The Y-axis represents percent inhibition compared to control and the X-axis represents the concentration of each respective protein in nM.

### 2.3 Production and Purification of recombinant Proteins in E. coli culture

We were unable to produce the wild type proteins in either insect or yeast cell cultures. Thus, we decided to produce a mutated form of PAP-S1 (PAP-S1R68G) with a native signal peptide. We believed this would stabilize protein production but would greatly reduce activity in *E. coli* and somewhat reduce activity in eukaryotic cells in *E. coli* cell culture [21, 24]. Native environment of both PAP-S1R68G (purity of 60%) and RTA/PAP-S1R68G (purity of 55%) production are shown in Figure 3.a (purity determined by GelAnalyzer2010). More than 400ug of each protein was produced. However, as shown in Figure 3.b, 6-His tag purification process was very inefficient and a large amount of protein was lost in the wash (kept for further analysis). This loss may have been due to the 6-His tag being hidden by the protein in its final conformation. It is possible the double band is due to protein degradation or unwanted recombination at the production site by *E. coli* cells.

**Figure 3.**
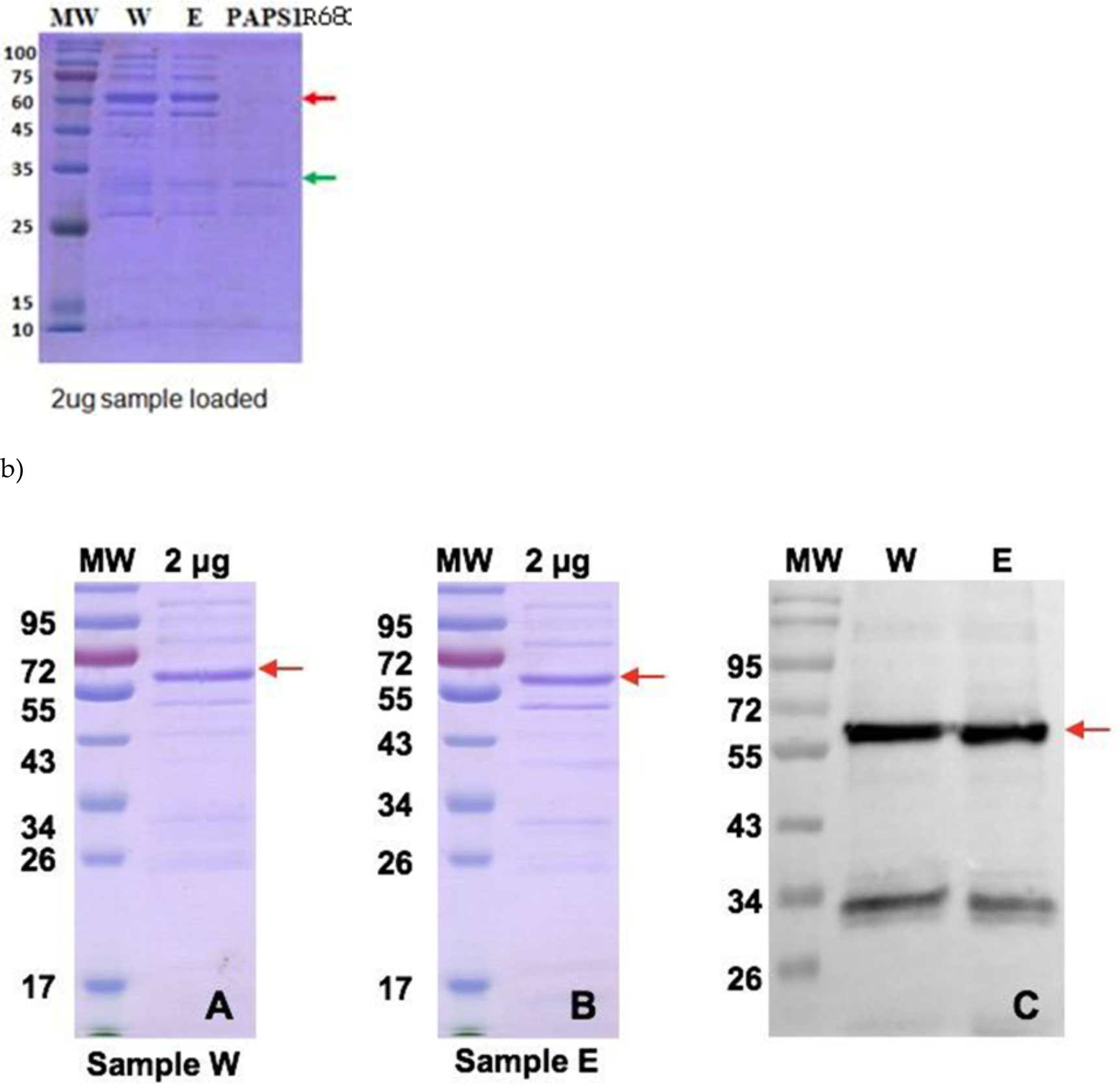
Protein Purification. a) Stained gel of RTA/PAP-S1R68G from third wash (sample W) and from pooled eluted fraction (sample E) at 60kDa (red arrow) and of PAP-S1R68G at 30kDa (green arrow), all from native environment, after buffer exchange. The purity of sample W was 65%, of sample E 55%, and of PAP-S1R68G 60%. b) Close up and Western Blot of sample W and E. A double band is present and may be due to degradation of the protein or as a result of an unwanted recombination by *E. coli* cells.

For each sample in Table 2, the final concentration was determined using a Bradford protein assay; purity was determined using the GelAnalyzer 2010 (see appendix 4 for details). The yield was very low, and again, a lot of protein was lost in the washes as the 6-His tag purification does not appear to be the right system for those proteins.

**Table 2.** Protein concentration and purity. For each sample, protein content was determined using a Bradford protein assay; purity was determined by GelAnalyzer 2010 as explained in appendix 4.

| Native protein | Final concentration | Volume | Final product | Purity |
| --- | --- | --- | --- | --- |
| PAP-S1R68G | 0.23mg/ml | 3.2ml | 0.736mg | 60% |
| RTA/PAP-S1R68G | 0.53mg/ml | 1.35ml | 0.716mg | 55% |
| (sample E) |  |  |  |  |
| RTA/PAP-S1R68G<br>(sample W) | 0.26mg/ml | 4.8ml | 1.25mg | 65% |

### 2.4 Inhibitory activity of recombinant proteins in the Rabbit Reticulate Lysate TnT^®^ system

The inhibitory activity of PAP-S1R68G and RTA/PAP-S1R68G were determined using 5 different concentrations of PAP-S1R68G and RTA/PAP-S1R68G (sample E) on the Rabbit Reticulate Lysate TnT^®^ system using Luciferase as control. A Luciferase assay was used to determine Luciferase expression levels using a luminometer. The comparative plot is shown in Figure 4 and includes previous data on Ricin and RTA obtained similarly [22]. As can be observed, RTA/PAP-S1R68G behaves more like RTA than PAP-S1R68G and has an IC50 at 0.025nM (similar to RTA 0.03nM) against 0.06nM for PAP-S1R68G. The total inhibition is attained at 0.83nM for RTA/PAP-S1R68G while PAP-S1R68G barely reaches 90% at 16.67nM, possibly due to the single point mutation (R68G). It is also interesting to note that PAP-S1R68G has about the same IC50 as PAP-S2 (0.07) but a much higher total inhibition point (around 1.2nM for PAP-S2) [2]. These results not only show that RTA/PAP-S1R68G is at least twice as fast as PAP-S1R58G but also 16 times more potent. It is comparable to native RTA and can thus be assumed that non-mutated RTA/PAP-S1 will be even faster.

**Figure 4.**
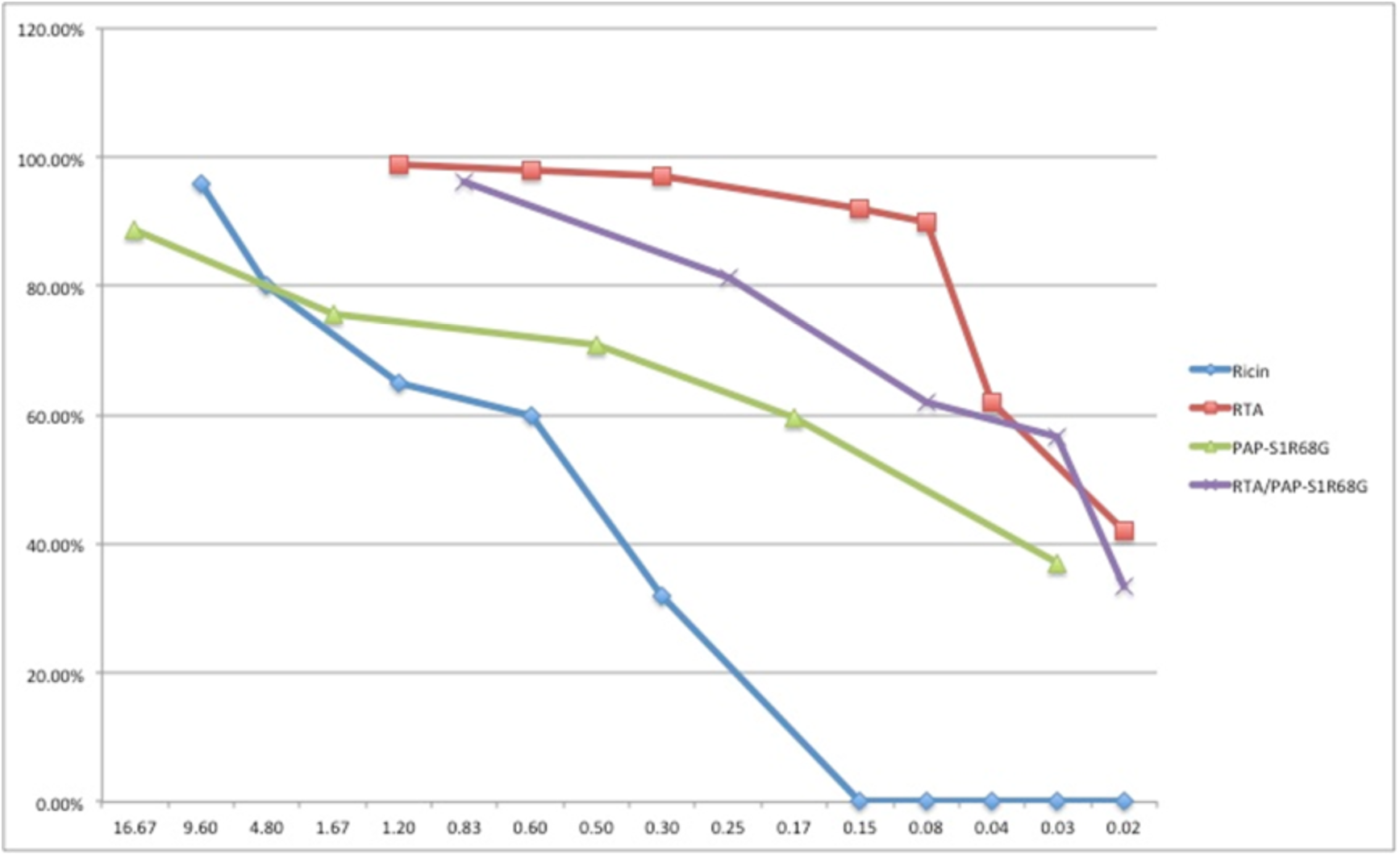
Comparative inhibitory activity. The Y-axis represents percent inhibition compared to control and the X-axis represents the concentration of each respective protein in nM. Results represent the average S.D. for two individual experiments.

The same inhibitory activity tests (Figure 5) were run under the same conditions for the proteins from sample W in order to determine whether binding or protein type isolation issues were causing the high amount of protein loss during washing. The results suggest behavior that is more like wild type Ricin protein, but with an IC50 of 0.3nM against 0.5nM for wild type Ricin.

**Figure 5.**
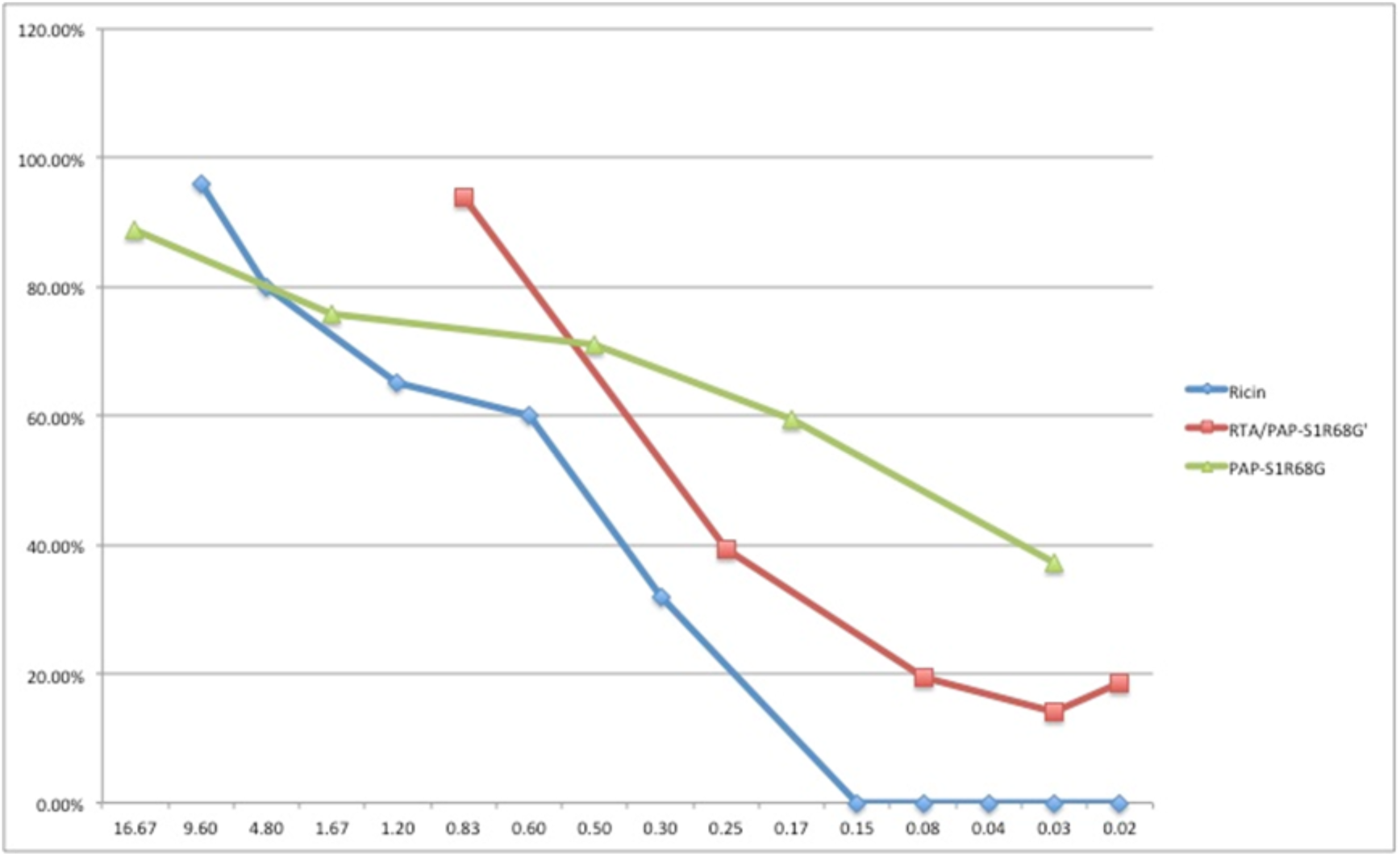
Comparative inhibitory activity. The Y-axis represents percent inhibition compared to control and the X-axis represents the concentration of each respective protein in nM. Results represent the average S.D. for two individual experiments.

The existence of two conformations of the same protein is possible as this fusion protein is new and it is difficult to predict exact conformation. Resequenced cDNA verified there were no acquired plasmid mutations. A prediction software [25] showed high probability (score of 0.9131) of existence of a di-sulfide bond between C173 and C539. This could explain the observed behavior and the difficulty in using 6-His tag for affinity purification. However, this remains to be proven, as this behavior was not observed in cell free expression system. Moreover, antibodies against PAP-S1 may solve the yield issue and any interaction with the 6-His tag itself.

## 3. Discussion

The fusion protein between Ricin A chain C terminus and PAP-S1 N terminus was observed to be functional and active in both eukaryotic and prokaryotic cell free systems with a great increase in both speed and potency compared to PAP-S1 alone. It was also observed that it was comparable in activity to Ricin A chain in a eukaryotic system. It is the opinion of the authors that additional research should be done in order to determine both the cytotoxicity and selectivity of RTA/PAP-S1 to PAP-S1 against a wide range of mammalian infectious diseases, including some types of cancers and a wide range of plant infectious diseases. We expect the RTA/PAP-S1 fusion protein to be a much more viable, potent, and less cytotoxic alternative to PAP-S1 alone for both agricultural and therapeutic applications.

## 4. Materials and Methods

The PureLink^®^ Genomic Plant DNA Purification Kit, HisPur™ Ni-NTA Spin Purification Kit, 0.2 m, Phire Plant Direct PCR Master Mix, PureLink™ Quick Gel Extraction and PCR Purification Combo Kit, Phire Hot Start II DNA Polymerase, Expressway™ Mini Cell-Free Expression System and NuPAGE™ 10% Bis-Tris Protein Gels, 1.5 mm, 15-well were purchased from Thermo Fisher Scientific. PCR Oligonucleotides and overlap extension products were synthesized by Integrated DNA Technologies. Rabbit Reticulate Lysate TnT^®^ Quick Coupled Transcription/Translation System, Luciferase Assay System and the *E. coli* S30 T7 High-Yield Protein Expression System were purchased from Promega. cDNA coding for the PAPS1[R68G] protein and Ricin-A-Chain/PAP-S1R68G was chemically synthesized with optimization for *E. coli* expression by GenScript. The *E. coli* pT7 expression vector and *E. coli* strains BL21(DE3) were purchased from Proteogenix. Total sample protein content was analyzed using Qubit™ 3.0 Fluorometer and Bradford protein assay. Luciferase assay readings were achieved using a Perkin Elmer EnVison Microplate Reader. Gel analyses were performed using a GelAnalyzer 2010.

### 4.1 E. coli cell free expression and E. coli protein synthesis inhibition

#### 4.1.1 Design of the DNA sequences of the proteins for *E. coli* cell free expression system

The DNA sequenced was designed and isolated as described in our previous work [20]. In short, fresh seeds of both *Ricinus communis* and *Phytolacca Americana* were purchased a local supplier in Baltimore, MD and Ricin A Chain (RTA) and PAP-S1 (with native signal peptide) isolated and PCR amplified. The 6-His tag was added to PAP-S1 at the C terminal. The RTA/PAP-S1 fusion protein was achieved through PCR extension, using the native RTA polylinker, between RTA C terminal and PAP-S1 (without the signal peptide) N terminal with the 6-His tag at RTA N terminal. The PAP-S1/RTA fusion protein was achieved through the same means but between PAP-S1 C terminal and RTA N terminal with the 6-His tag at PAP-S1 N terminal.

#### 4.1.2 *E. coli* cell free expression and purification

The PAP-S1, PAP-S1/RTA and RTA/PAP-S1 fusion proteins were produced using Expressway™ Mini Cell-Free Expression System as previously described [20]. In short, Linear DNA was used for all proteins in thrice the volume (150uL) for PAP-S1 and PAP-S1/RTA and twice the volume for RTA/PAP-S1 using the T7 promoter System. The proteins were purified using the HisPur™ Ni-NTA Spin Purification Kit, 0.2 m before being run on protein gels for confirmation. The total protein content was determined using Qubit™ 3.0 Fluorometer. Gels were analyzed using the GelAnalyzer2010.

#### 4.1.3 Inhibition of protein synthesis in cell free *E. coli* system

Enzyme activity of the purified recombinant proteins was determined by the intensity of the band on protein gel of a control against expression of the control without the recombinant proteins, after protein purification using the HisPur™ Ni-NTA Spin Purification Kit, 0.2 m, as previously described [20] using the *E. coli* S30 T7 High-Yield Protein Expression System. The control used was the pEXP5-NT/CALML3 control vector with a DNA template expressing an N-terminally-tagged human calmodulin-like 3 (CALML3) protein (under the T7 promoter). The concentration of CALML3 was determined for increasing concentrations of recombinant PAP-S1 and fusion proteins by measuring band intensity on a protein gel (Coomassie blue stained) by GelAnalyzer2010.

### 4.2 E. coli in vivo expression system and Rabbit Reticulate Lysate protein synthesis inhibition

#### 4.2.1 Design of the DNA sequences of the proteins for *E. coli in vivo* expression system

The cDNA coding for a mutated version of PAP-S1 and RTA/PAP-S1, namely PAP-S1R68G and RTA/PAP-S1R68G, were chemically synthesized with optimization for *E. coli* expression by GenScript. The mutated form of PAP-S1 was used in order to reduce *E. coli* ribosome depurination by PAP-S1 and RTA/PAP-S1while safeguarding their Eukaryotic ribosome depurination activities [21]. The native PAP-S1 signal peptide was kept for PAP-S1R68G with the addition of an *E. coli* signal peptide of two amino acids at PAP-S1R68G and RTA/PAP-S1R68G N terminal and with the 6-His tag at PAP-S1R68G and RTA/PAP-S1R68G C terminal (see appendix 1 for details).

#### 4.2.2 *E. coli in vivo* expression vector

The cDNA sequences described above were cloned in an *E. coli* pT7 expression vector using the Ncol/XhoL1 cloning strategy (map of vectors in Appendix 1).

#### 4.2.3 *E. coli* protein production

Optimal conditions for *E. coli* Bl21(DE3) and modified Bl21(DE3) were determined for PAP-S1R68G and RTA/PAP-S1R68G respectively in small volumes before being scaled up to 1L production culture (please contact the authors for more details). In short, bacteria starter were obtained by incubation at 37°C and then followed by IPTG induction at specific temperatures and incubation times. The bacteria were then harvested by centrifugation, followed by lysis. The supernatant was collected after centrifugation for both proteins, the native proteins extracts.

#### 4.2.4 *E. coli* protein purification

The purification of the native proteins extracts was achieved by affinity versus His-tag on Ni-resin. The equilibration was done with a standard binding buffer. Wash and elution was performed by imidazole shift. After purification, fractions of interest were pooled, concentrated, and analyzed by SDS-PAGE. The final concentration was determined by Bradford protein assay.

#### 4.2.5 Rabbit Reticulate Lysate protein synthesis inhibition

The inhibitory activity of PAP-S1[R68G] and RTA-PAP-S1[R68G] were tested by using the Rabbit Reticulate Lysate TnT^®^ Quick Coupled Transcription/Translation System and the Luciferase Assay System. Briefly, each transcription/translation reaction run was performed according to the instructions for use (IFU) in the presence of a T7 Luciferase reporter DNA, and the Luciferase expression level was determined with a Perkin Elmer EnVison Microplate Reader. Transcription/translation runs were done twice with and without addition of five different concentrations of PAP-S1R68G and RTA-PAP-S1R68G in order to determine the inhibitory effect of the proteins. PAP-S1R68G and RTA-PAP-S1R68G concentrations were adjusted by taking sample purity into consideration.

